# Evidence of predictive information compression in latent space in humans during speech listening

**DOI:** 10.64898/2026.07.14.738305

**Authors:** A. Corsini, S. Schneider, A. Tomassini, L. Pedani, L. Fadiga, A. D’Ausilio

**Affiliations:** Center for Translational Neurophysiology of Speech and Communication, Istituto Italiano di Tecnologia, Ferrara, Italy; Department of Neuroscience and Rehabilitation, Università di Ferrara, Ferrara, Italy; Institute of Computational Biology, Computational Health Center, Helmholtz Munich, Munich, Germany

## Abstract

Speech perception requires transforming acoustic input into neural representations that support linguistic understanding, yet its underlying computational principles remain unclear. Classical efficient coding theories posit optimal compression of sensory input, whereas alternative accounts propose that neural systems preferentially encode information that supports prediction. A key open question is whether such predictive encoding operates on fixed inputs or on flexible internal representations. We instantiated three hypothesis models of speech processing: (i) optimal compression with deep autoencoders, (ii) predictive reconstruction with predictive autoencoders, and (iii) predictive information representation via latent-space prediction using contrastive learning. We compared resulting speech latent representations to electroencephalographic (EEG) activity during speech listening. Representations learned under the predictive information objective best explained neural latents. Crucially, only representations that selectively compressed predictive information predicted behavioral performance, suggesting that neural speech representations are structured to encode predictive information in latent space rather than to maximize compression or input prediction.

## Main

The ultimate goal of perception is *explanation*—the ability to infer representations of the world from sensory inputs that are useful for prediction and behavior.^1^ As an example, speech perception is a task in which successful comprehension requires extracting relevant acoustic information from signals in which informative and irrelevant components are nonlinearly mixed^2^. Classic sensory coding theories argue for the importance of information maximization about the input stimulus under channel capacity constraints. ^3–6^ Alternatively, the nervous system may follow a predictive encoding principle, producing efficient representations of predictive information. ^7–9^ Although both principles implement forms of efficient coding, they differ fundamentally: the former lacks an explicit notion of relevant information within signal variability and therefore assumes that no information should be discarded, whereas the latter selectively prioritizes information that supports prediction.

Prediction is a fundamental function of the brain for guiding future actions. For instance, when catching a ball, sensory inputs must inform future movements that are inherently delayed due to biological constraints such as sensory transduction and muscle activation^10^. From another perspective, actions can be viewed as means to reach an expected sensory state^11,12^, such as a proprioceptive state indicating that the ball has been successfully caught. In both cases, a representation of the trajectory of the ball is useful only insofar as it allows predicting its evolution sufficiently far into the future. This idea that the neural system should learn an efficient ‘*small-scale model’* from which to generate predictions dates back to Craik *(1943)*, who first proposed such predictions should occur in a *learnable* symbolic (latent) space. However, whether and how these theoretical principles map onto computations during speech perception remains largely unknown.

M/EEG studies have often circumvented this problem by focusing on specific, *a priori* features of the speech signal known to drive cortical activity, such as the envelope in delta/theta bands^13–16^ spectrograms^17^ or phonetic features^18^. While successful in linking neural activity to specific features of speech, they a priori constrain the representational space and therefore offer limited insight into how the brain autonomously extracts relevant information from the full sensory input. More recently, deep learning models developed for automatic speech recognition (ASR),^19,20^ and text generation (LLMs) have been used to generate rich, high-dimensional representations of speech and language. These representations have enabled hypothesis testing about the nature of internal representations during naturalistic listening ^21,22 23–26^. For example, Caucheteux and colleagues *(2022)*^*27*^ used the deep language model GPT-2 to show how the brain makes simultaneous predictions at different hierarchical levels of the linguistic structure, from frontoparietal to temporal cortices.

However, none of these approaches explicitly tests how the neural system constructs efficient internal models of speech. Specifically, during speech listening, neural activity could be optimized to follow (i) an information-maximization principle to produce efficient codes of the speech stimulus via optimal compression, (ii) a predictive reconstruction principle through prediction error minimization about the future speech input, or (iii) a predictive information principle via prediction in a learnable latent space. To test these hypotheses, we first extract task-relevant information using a state-of-the-art speech recognition model (wav2vec 2.0), thereby operating in a high-performance speech representation space. We then instantiate these principles into alternative deep neural networks (DNNs) by modifying their architecture and objective function. We train these models and compare their latent speech representations with EEG activity of 26 participants listening to the same sentences while performing a speech recognition task (Fig 1a).

**Figure 1.**
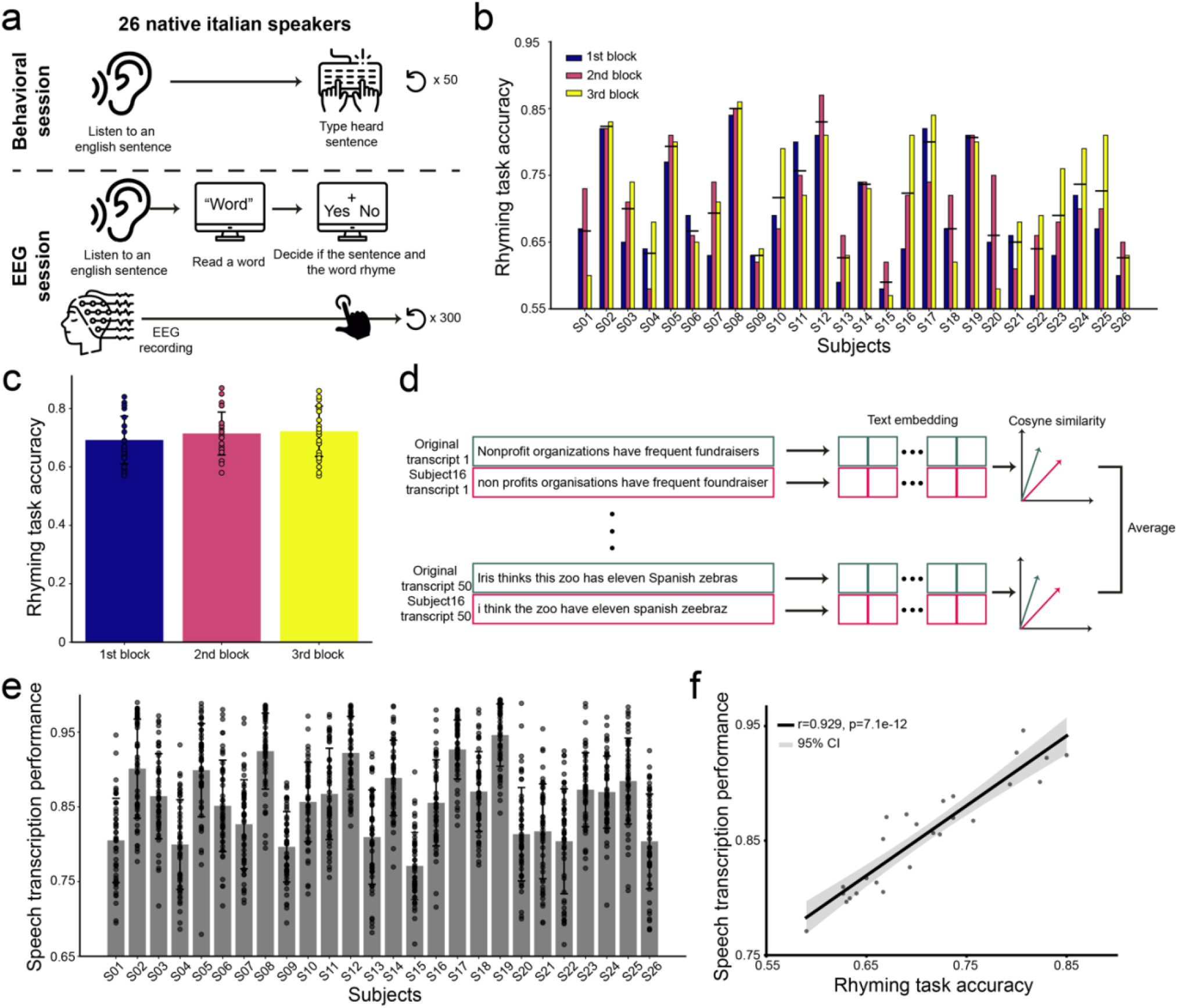
Experiment design and behavioral results. **a**. Overview of the experimental sessions. Top: speech transcription task performed outside the EEG session; bottom: EEG session with a rhyming task following each sentence. **b**. Individual accuracies in the rhyming task across the three EEG blocks (n = 26 participants, 300 trials each, 100 per block). **c**. Mean rhyming task accuracy did not differ significantly across blocks, confirming performance stability. **d**. Extraction of the speech transcription performance from the behavioral session. Participants transcribed half of the English sentences presented in the EEG session; transcripts were compared to ground-truth sentences using text-embedding cosine similarity. **e**. Distribution of transcription accuracies across participants. **f**. Rhyming task accuracy strongly correlated with individual transcription accuracies (*r* = 0.929, *p* = 7.1 × 10^-12^), indicating that the rhyming task provides a reliable behavioral proxy for active neural encoding of speech related to speech recognition.

To implement the proposed hypotheses in DNN models we trained (i) a deep autoencoder (deep-AE) that learns optimal nonlinear compression of speech features using a decoder trained with a reconstruction loss, (ii) a deep predictive autoencoder (deep-pAE) that learns optimal nonlinear compression of speech features with a decoder that minimizes the prediction error about the future input and (iii) a contrastive learning model that learns a representation of the predictive information by optimizing a loss in the latent space with no reconstruction (CL)^*28*^. Among these models, CL representations best explained neural activity. Furthermore, when CL was trained to compress predictive information using an output bottleneck, neural encoding of resulting speech representations correlated with participants’ speech recognition performance. This selective alignment suggests that a key principle of neural information processing during speech perception is the extraction of predictive information through prediction in latent space, enabling the system to discard irrelevant signal details. Beyond providing insight into the neural computations underlying speech perception, our results offer a principled framework for designing DNN-based hypotheses of speech perception whose representations can be directly compared with neural latent representations measured in human EEG.

## Results

### Speech recognition performance

We analyzed the EEG activity of 26 healthy Italian participants listening to short English sentences (n=100; from 2.9 to 4.5 seconds duration). All participants were native Italian speakers who self-reported English as their second language (the use of a second language ensures greater intersubject variability and avoids ceiling effects in comprehension performances). Stimuli were selected from the USC-Timit dataset^29^ which provides recordings of American English speakers, along with their transcripts. Each sentence was presented three times in a randomized order during the experiment, resulting in a total of 300 trials per participant, divided in three blocks. Each sentence was followed by a speech recognition task (Fig. 1a bottom). The task consisted in the visual presentation of one word at the end of the played audio which could either rhyme or not rhyme with any of the words in the heard sentence. This task implicitly forces participants to perform an internal transcription of the words to be compared with the one presented at screen. Participants were asked to indicate whether the word rhymed or not by pressing a button. To avoid response bias, rhyming and non-rhyming words were matched for their average frequency of use (~6e-05). Participants’ success rate in the blocks ranged from 57% to 87% (Fig 1b). Average success rate in the three blocks was not significantly different (Fig. 1c), thus ensuring compatible results across the whole session. To verify that the designed rhyming task is a good proxy of speech-to-text skills, we assessed participants’ speech transcription skills in a separate behavioral session (Fig. 1a top). They listened to half of the same English sentences, and at the end of each one, they were asked to transcribe them as accurately as possible. The transcriptions were then compared to the true sentences using OpenAI’s text embedding model, text-embedding-ada-002. Specifically, each participant’s transcript was converted into an embedding vector, and its cosine similarity with the embedding vector of the correct sentence was computed (Fig. 1d). This similarity was then averaged across sentences for each participant, serving as a score for their speech transcription performance (Fig. 1e). Behavioral performance in rhyming task was strongly correlated to speech transcription performance (*r* = 0.929, *p* = 7.11e-12; Fig. 1f), suggesting that it can serve as a reliable proxy for the active neural encoding of speech underlying speech recognition.

### Neural encoding of wav2vec 2.0 features does not correlate with participants’ speech recognition performance

We first transformed speech signals using the wav2vec 2.0 (Extended Data Fig. 1a), which is the current state-of-the-art for speech to text tasks^19^. Features extracted by wav2vec 2.0 are expected to retain primarily the information necessary to perform speech recognition. Moreover, wav2vec 2.0 features are more effective when decoding M/EEG signals than simple spectrogram features (Mel Spectrogram) and features learnt end-to-end together with M/EEG data (Deep Mel)^30^.

To check that wav2vec 2.0 features are relevant for participants during speech listening, we computed mutual information (MI) between the participants’ EEG activity and the output wav2vec 2.0 latents of the same speech signals using the Gaussian Copula Mutual Information (GCMI) estimator^31^. We expected this to be encoded in the listeners’ activity with a reasonable lag and to show a meaningful topography of information encoding. At the same time, we hypothesized that the strength of this encoding would not predict participants’ behavioral performance in the speech recognition task. Indeed, unlike biological systems, wav2vec 2.0 is optimized solely for recognition performance, whereas neural representations are shaped by efficiency constraints that prioritize task-relevant information under biological limitations.

In line with previous findings^22,25^, we found that wav2vec 2.0 features are significantly encoded in the listeners’ neural activity (*p* = 2.00e-04; one side cluster-based statistics^32^ against the mean of 100 surrogate data; Extended Data Fig 1b). This encoding was primarily localized over central and temporal electrodes (Extended Data Fig 1b). However, in line with our hypothesis, the strength of encoding of the features did not correlate with participants’ speech recognition performance (*r* = 0.09, *p* = 6.44e-01; Extended Data Fig. 1c).

To our knowledge, this is the first evidence of such a dissociation: although wav2vec 2.0 representations are significantly encoded in neural activity, they do not appear to isolate the subset of information that is behaviorally relevant for human speech recognition. Together, these results indicate that wav2vec 2.0 features provide a strong and neurally grounded representational basis yet remain insufficient to fully capture the computations underlying human behavior.

### Linear dimensionality reduction does not isolate relevant information

To make sure that linear dimensionality reduction cannot fill the missing correlation with behavior, we ran a principal component analysis (PCA)^33^, that, beyond visualization, is often employed in neuroscience to reduce dimensionality while preserving relevant information^34–36^. In a similar way, we, here, try to isolate the behaviorally relevant information in the dataset using PCA on wav2vec

2.0 features. Because PCA orders components by explained variance, components with small eigenvalues are often treated as noise and discarded. However, this heuristic can obscure structured, task-relevant information that resides in low-variance dimensions, for example, when solving image recognition with deep neural networks^37^.

For this reason, we applied PCA to the wav2vec 2.0 output and separated it into top- and bottom-variance subspaces (Fig. 2b, left). Top variance subspaces are obtained by cumulating explained variance ratio starting from the most informative feature (PC1) to reach a targeted value, conversely bottom variance subspaces are generated to reach a targeted explained variance value starting from the least informative feature (the last PC; Fig. 2a). Even subspaces explaining as little as 0.1% of variance retained time-specific information and spectral content consistent with top subspaces (Fig. 2b, right). Sentence correlation matrices revealed lower correlations in bottom subspaces, suggesting they encode more sentence-specific information (Fig. 2c).

**Figure 2.**
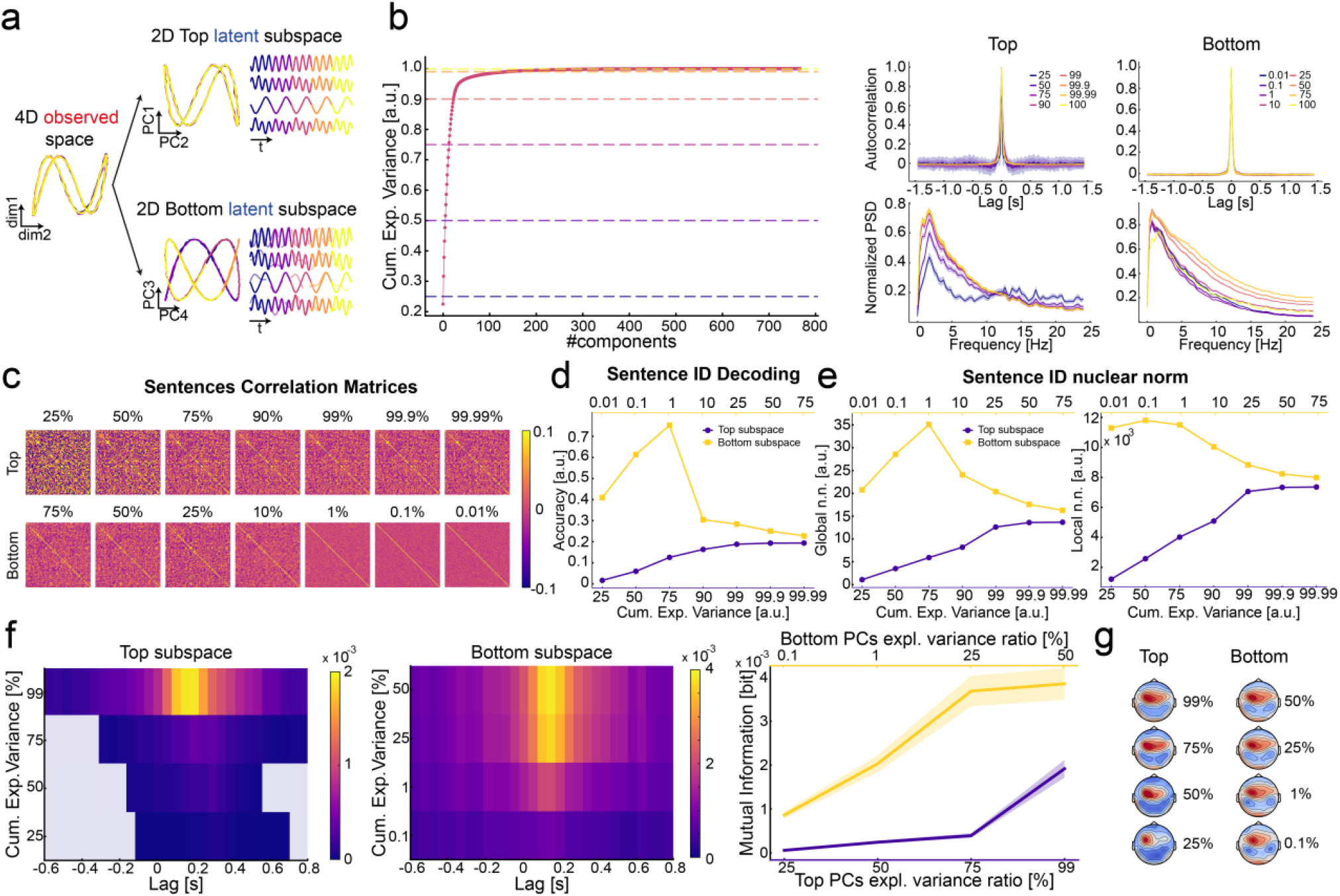
Linear dimensionality reduction does not isolate relevant information. **a**. PCA on a synthetic 4D dataset. The first two PCs explain >99% of the variance and capture the main periodic structure, whereas the last two PCs, though low in variance, more clearly disentangle the periodic components, while being bad at signal reconstruction. **b**. PCA of wav2vec 2.0 features from the speech dataset used in the EEG experiment. Left: cumulative explained variance of the 768 PCs. Eight subspaces were selected based on explained variance, from both the top (25–100%) and bottom (0.01–100%) variance ranges. Right: autocorrelation and normalized power spectral density (PSD) show strong temporal specificity across all subspaces. **c**. Correlation matrices of the 100 sentences represented in each subspace. Sentence correlations are higher for top-variance subspaces, indicating less sentence-specific structure. **d**. Sentence ID decoding accuracy using a kNN classifier. Performance is low for top subspaces and saturates near 99% variance, whereas bottom subspaces perform better, peaking at 75.3% for the 1% variance subspace. **e**. Global and local nuclear norms of PCA subspaces (sentences as classes). Bottom subspaces show higher global nuclear norm (greater class separability) and higher local nuclear norm (richer, less compressed sentence representations). **f**. Time-lagged mutual information (MI) between speech PCA subspaces and listeners’ EEG activity. Bottom subspaces show higher MI than top ones, peaking at a physiologically consistent 0.15 s lag. EEG encodes as much information from the bottom 1% subspace as from the top 99%. **g**. MI topographies at peak lag. All subspaces, including low-variance ones, show plausible central distributions.

Decoding analyses confirmed this: when retaining increasingly large fractions of variance in the top subspaces (25 – 99.99 %), decoding accuracy improved up to ~90% explained variance, after which gains plateaued (Fig. 2d). In contrast, when high-variance components were excluded, decoding accuracy peaked in the bottom 1% variance subspace (75.3%), indicating that low-variance PCs carry meaningful information often overlooked by standard PCA practice. Global and local nuclear norm analyses supported these results. In practice, global nuclear norm is a proxy of linear separability of the classes (the higher the better), while local nuclear norm is a measure of the compactness of the data (the higher the less compact). Bottom subspaces exhibited higher global separability (lower rank between classes) and greater local complexity (higher rank within sentences) (Fig. 2e).

To test whether these PCA subspaces are encoded in the neural activity and, at the same time, correlate with participants’ speech recognition performance, we computed MI between PCA subspaces and EEG signals. EEG activity predominantly encoded bottom-variance subspaces: the bottom 1% explained variance shared the same amount of information with EEG (0.0020 ± 0.00019 bits) than the remaining 99% (0.0019 ± 0.0002 bits; Fig. 2f, right). Even subspaces accounting for the 0.1% of variance showed a coherent peak of information encoding (0.15s) and a meaningful topographic distribution of the information (Fig. 2g). However, most importantly, we did not find any significant correlation between EEG encoding of any of the selected top- (25%: *r* = 0.201, *p* = 3.22e-01; 50%: *r* = 0.162, *p* = 4.28e-01; 75%: *r* = 0.134, *p* = 5.12e-01; 99%: *r* = 0.209, *p* = 3.03e-01; Extended Data Fig. 2a) and bottom-variance (0.1%: *r* = 0.093, *p* = 6.48e-01; 1%: *r* = 0.084, *p* = 6.80e-01; 25%: *r* = 0.144, *p* = 4.80e-01; 50%: *r* = 0.141, *p* = 4.90e-01, Extended Data Fig. 2b) subspaces with participants’ speech recognition performance.

Together, these results show that even if both high and low variance components of wav2vec 2.0 feature are significantly encoded in listeners’ EEG activity, the encoding strength of none of the selected subspaces correlates with their speech recognition performance.

### Predictive information is best encoded in neural activity and its compression explains participants’ speech recognition performance

We now want to test alternative encoding hypotheses on speech perception. To do so we need to move beyond PCA and constrain speech features using more complex nonlinear models. To our knowledge, no previous work has tried to test speech perception hypotheses by changing the architecture and the objective function of DNN models and comparing the resulting speech latents with human neural latents (Fig. 3a-b)

**Figure 3.**
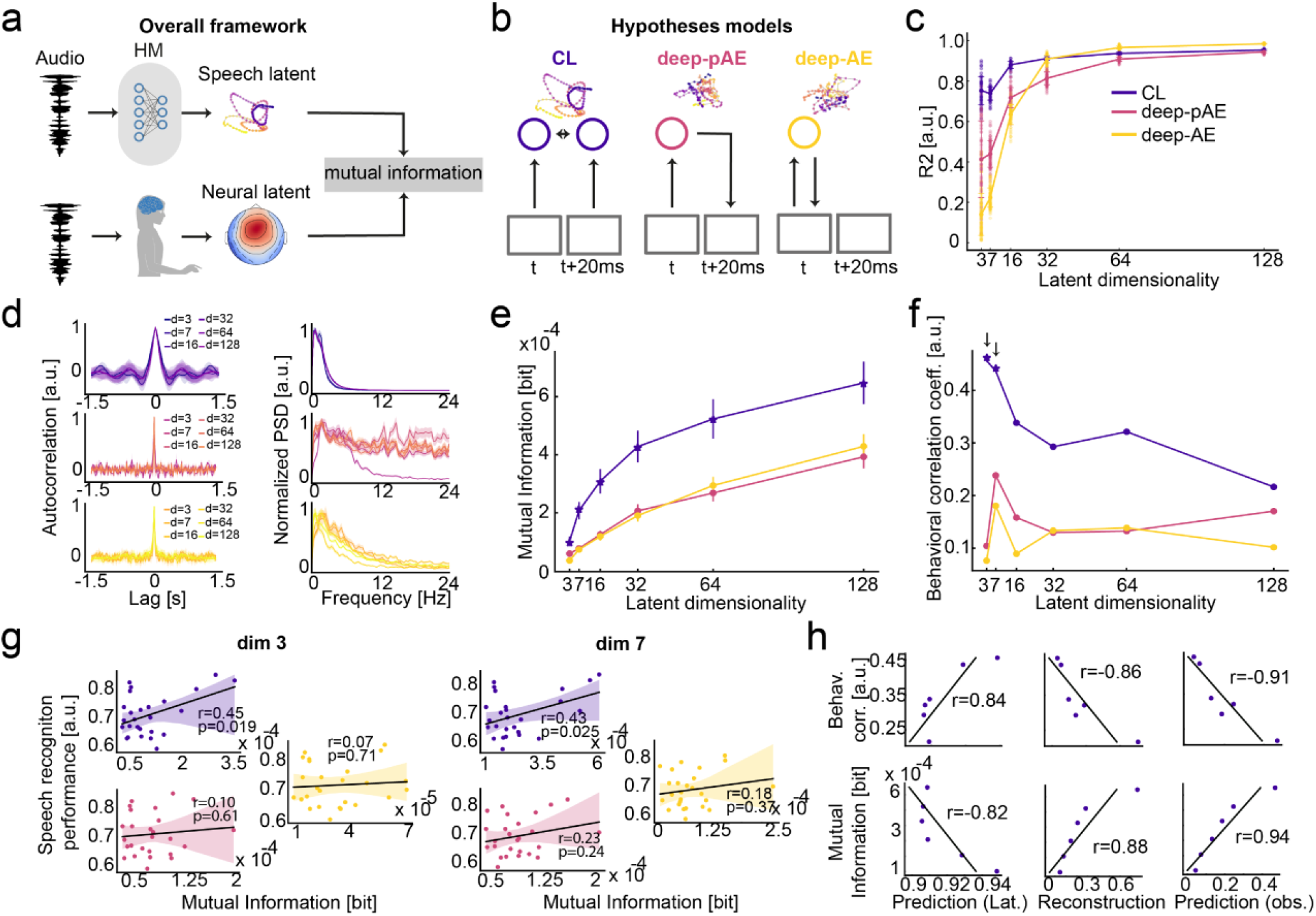
Predictive information is best encoded in neural activity and its compression explains speech recognition performance. **a**. Model–brain comparison schematic. Speech features were encoded by each trained model into speech latents, while 26 participants listened to the same stimuli during EEG recording (neural latents); mutual information (MI) was computed between the two. **b**. Three speech feature reduction architectures. Contrastive learning (CL) enforces temporal predictability between consecutive embeddings; the predictive autoencoder (deep-pAE) learns latent states forecasting future signal evolution; the standard autoencoder (deep-AE) compresses features for present-signal reconstruction. **c**. Embedding consistency across training runs. CL yields highly consistent representations even at very low latent dimensionalities. **d**. Temporal structure of latent features. Autocorrelation (left) reveals sustained predictability only for CL; power spectra (right) show CL consistently extracts low-frequency components across dimensionalities. **e**. MI between neural and speech latents across output dimensionalities. CL features share more information with EEG than both autoencoder models. **f**. Across-subject correlations between MI and speech recognition performance. Only CL shows consistently strong correlations, reaching significance at dimensionalities 3 and 7. **g**. Scatter plots of MI versus speech recognition performance at the two significant dimensionalities (3 and 7). **h**. CL models excelling at latent prediction (lower dimensionalities) share more behaviorally relevant information with EEG; the reverse holds for models excelling at signal reconstruction.

We first verified that our three models produce sufficiently distinct speech latents reflecting the initial hypotheses: (i) optimal compression via a deep autoencoder (deep-AE) optimizing nonlinear reconstruction of wav2vec 2.0 features, (ii) predictive information in signal space with a predictive deep autoencoder (deep-pAE) minimizing future prediction error, and (iii) predictive information in latent space using a contrastive learning model (CL) optimizing latent-space prediction via InfoNCE loss (Extended Data Fig. 3a–b). CL representations showed the highest consistency across training runs, remaining reproducible even at very low dimensionalities^28^ (Fig. 3c), and exhibited strong temporal predictability with consistent spectral structure across dimensions (Fig. 3d). Deep-AE and deep-pAE performed well on reconstruction but showed limited predictive accuracy at low dimensionality, whereas CL supported superior temporal prediction even with three latent dimensions, consistent with compact spaces favoring isolation of predictive information. Dimensionality seven was chosen as an effective bottleneck, above which predictive information extraction improves by no more than 1% (Extended Data Fig. 3c). CL representations also remained more temporally predictive than deep-pAE latents at larger offsets, despite no explicit multi-step training (Extended Data Fig. 4b–d).

To determine which hypothesis best accounts for neural computations, we quantified EEG encoding of each model’s speech latents and tested whether encoding strength correlated with speech recognition performance. The best-fitting hypothesis should (i) yield the most strongly neurally encoded features and (ii) show a positive encoding strength–performance correlation.

To this end, we computed mutual information between speech latents and neural latents (Fig. 3a). Across all output dimensionalities, CL latents were significantly more strongly encoded in the EEG than both autoencoder models (Fig. 3e; CL vs p-AE: *p* = 1.81e-02, *p* = 3.04e-05, *p* = 1.57e-05, *p* = 9.71e-06, *p* = 2.09e-05, *p* = 2.92e-06; CL vs AE: *p* = 1.18e-03, *p* = 1.34e-05, *p* = 1.49e-05, *p* = 6.06e-06, *p* = 6.53e-05, *p* = 1.20e-05; t-test for related samples, Bonferroni corrected). This advantage persisted in time-resolved analyses (Extended Data Fig. 5a), where CL consistently showed broader and earlier windows of significant neural encoding than both AEs across output dimensions. Therefore, producing speech representations through predictive information maximization via prediction in latent space better explained neural activity. Crucially, only the average encoding of low-dimensional CL latents correlated positively with speech recognition performance (dim 3: *r* = 0.45, *p* = 1.91e–02; dim 7: *r* = 0.43, *p* = 2.56e–02; Fig. 3f-g). In fact, CL models that emphasized signal reconstruction explained less behavioral variance, while those compressing predictive information in latent space performed systematically better (Fig. 3h). The temporal profile of these correlations mirrored the neural encoding pattern, peaking at slightly earlier lags (Extended Data Fig. 5b). Therefore, only compression of predictive information in latent space satisfies the original hypotheses.

To further test the alignment between the CL speech latents and neural latents, we computed MI between CL latents and PCA subspaces of wav2vec 2.0. CL preferentially represented low-variance components, with the bottom 25% variance subspace contributing much more (6.24 bits) than the top 75% (1.52 bits; Extended Data Fig. 5c). This pattern became stronger when CL was trained to predict further into the future (Extended Data Fig. 5c).

Overall, the results are compatible with the hypothesis that neural latents are optimized to represent predictive information, and that speech recognition performance are driven by the compressed representation of this predictive information.

### Speech neural encoding parallels non redundant predictive information

If the brain follows a similar predictive encoding strategy, we should observe a comparable temporal pattern of predictive information accumulation in both speech and neural latents. To test this, we compared how the CL model accumulates predictive speech information at inference time (Fig. 4a) with how neural latents encode predictive information during speech listening. Although the model is trained on next-sample prediction (20 ms ahead), its predictive and memorization abilities are evaluated over a larger window spanning 600 ms before to 800 ms after speech sample presentation, consistent with previous EEG encoding analyses. A dominant temporal regularity in speech, and a prominent feature of EEG speech tracking, is the syllable, whose duration across languages, including English, is approximately 200 ms. We therefore examined the accumulation of predictive information (ΔI) in the 200 ms preceding speech sample presentation (Fig. 4b). At each 20 ms step prior to presentation, the model acquires predictive information about future speech beyond what was already accumulated (δi, Fig. 4a), a quantity we term *non-redundant predictive information*, as it is, to a first-order approximation, predictive information not redundant with that acquired in the past.

**Figure 4.**
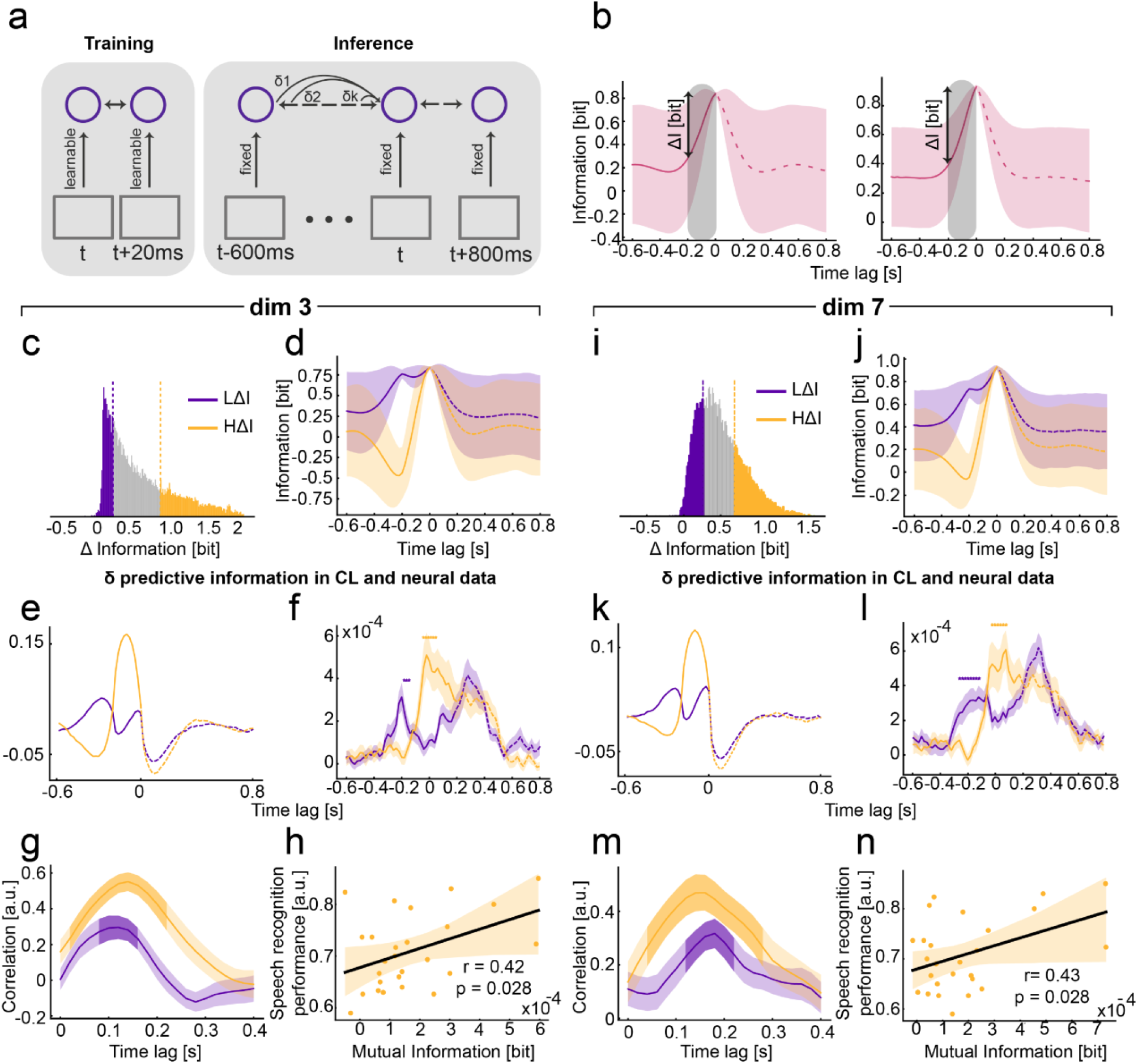
Speech neural encoding parallels non-redundant predictive information. **a**. Analysis schematic. The CL model is trained on next-sample prediction in latent space (20 ms) and evaluated over a wider inference window (−600 to +800 ms). At each speech sample prior to a target at time *t*, the model accumulates additional non-redundant predictive information (δ-information) about that target. **b**. Inference-time per-sample CL loss across the full window for latent dimensionalities 3 (left) and 7 (right), reflecting information about a given sample contributed by both preceding (prediction) and succeeding (memorization) context. **c–d**. (Dim 3) Distribution of per-sample CL loss when predicting 200 ms ahead (c), used to split samples into high- (HΔI) and low-(LΔI) predictive information classes based solely on model-derived measures. Inference-time loss curves for the two classes are shown in (d). **e**. (Dim 3) δ-information for HΔI and LΔI samples: the non-redundant predictive information extracted at each step is greater for HΔI samples, reflecting their higher accumulation demand. **f**. (Dim 3) EEG encoding for HΔI and LΔI samples. Neural encoding tracks the non-redundant information accumulated in the 200 ms before stimulus onset. **g**. (Dim 3) Cross-correlation between δ-information and EEG encoding peaks at a biologically plausible delay of 140 ms. **h**. (Dim 3). Only EEG information corresponding to high-ΔI samples within the key latency windows (< 140 ms) correlates with participants’ speech recognition performance (*r* = 0.42, *p* = 2.8e-02). **i–n**. (Dim 7) Panels i–n replicate panels c–h for latent dimensionality 7, yielding consistent results and conclusions.

We split speech samples into two classes (Fig. 4c, 4i): those easily predictable 200 ms in advance (low-Δ-information: LΔI) and those that are not (high-Δ-information: HΔI) (Fig. 4d, 4j). Importantly, the two classes do not differ in predictability at the moment of presentation, the sole distinction lies in how much predictive information was accumulated beforehand. Non-redundant predictive information (δ-information), defined as the temporal derivative of ΔI, was consistently higher for HΔI samples during this window, reflecting the greater accumulation required to compensate for their initially lower ΔI (Fig. 4e, 4k).

EEG encoding computed separately for HΔI and LΔI samples revealed the expected parallel: HΔI neural encoding was initially lower but rose rapidly to exceed LΔI within ~100 ms of onset (Fig. 4f, 4l; dim 3: *p* = 4.00e–02, 3.40e–03; dim 7: *p* = 1.60e–03, 3.44e–02), and the cross-correlation between δ-information and EEG encoding peaked at biologically plausible delays (Fig. 4g, 4m). Crucially, only the neural encoding of HΔI samples during the early information-gathering window correlated with behavioral performance (dim 3: r = 0.428, p = 2.89e–02; dim 7: r = 0.430, p = 2.81e– 02; Fig. 4f, right), whereas later or LΔI encodings did not. Together, these results suggest that behaviorally relevant neural responses are shaped by the early extraction of non-redundant predictive information.

## Discussion

We show that neural activity during speech perception is best explained by a system optimized for the extraction of predictive information. Speech representations learned under this principle were consistently better encoded in neural activity than those derived from models based on optimal information compression. The central role of prediction is not surprising, given its ubiquity across brain functions and the well-established presence of predictive computations in speech processing at multiple levels, from phonemes and syllables to words^39–41^, and even across longer temporal scales^27^. However, rather than focusing on prediction during speech inference, we approached the problem from a complementary perspective: how the brain learns the internal speech representations that are later used to support perception. In principle, different learning strategies could underlie speech perception. For example, the brain might first learn optimally compressed representations of speech, as proposed in classic efficient coding theories^4,5^, and subsequently build predictive mechanisms on top of these representations to guide perception. Under this hypothesis, models optimized for information compression should best explain neural activity, and their representations should be encoded in neural signals prior to stimulus onset (i.e., at negative lags), reflecting anticipatory predictions. However, our results do not support this scenario. Instead, models trained to extract predictive information produced representations that were more strongly encoded in neural activity, both at negative and positive temporal lags. This suggests that such representations are prioritized both by anticipatory processes (negative lags) and by stimulus-driven computations (positive lags) during speech perception. Moreover, the temporal accumulation of speech information in neural activity closely tracked the model’s extraction of non-redundant predictive information within a critical ~200 ms speech window. Together, these findings support the idea that the brain relies on a predictive information principle during speech listening and learning.

Our results, further show that speech perception is driven by the *compression* of such predictive information. Indeed, only low-dimensional output spaces produced speech representations whose neural encoding positively correlated with participants’ speech recognition performance. Using the InfoNCE objective, we identified the minimal latent dimensionality that maximized predictive information and observed that higher-dimensional representations increasingly incorporated reconstructive (and thus less task-relevant) features. These findings are consistent with the notion that the nervous system operates under metabolic constraints^42^ and that efficient information representation may confer evolutionary advantage^43^. However, while previous findings have highlighted the importance of compressing predictive information in the visual system^8,44–46^ and in artificial representation learning models^47,48^, it has not, to our knowledge, been directly examined in speech perception. Importantly, although compressed predictive representations best explained behavioral performance, overall neural activity was better captured by higher-dimensional, redundant representations. The role of redundancy in neural systems has long been debated^49^, and complete redundancy reduction is now considered neither achievable nor necessarily desirable^50–52^ particularly given its benefits for robustness and error correction. Rather than eliminating redundancy, the nervous system may instead represent and segregate redundant and non-redundant components in an explicit and functionally accessible manner^49^. In line with this view, our results suggest that while redundant, high-dimensional representations account for global neural activity, it is the explicit, capacity-limited representation of non-redundant predictive information that drives behavioral performance.

Finally, we found that such representations of compressed predictive information emerge specifically through prediction in a learnable latent space. The convenience of learning in an abstract, symbolic space was first proposed by Kenneth Craik (1943)^1^ who first suggested that the brain constructs internal models of the world by performing predictions in a latent representational space. This idea has since been formalized in hierarchical predictive coding frameworks.^53–56^ While not strictly a predictive coding model, our best contrastive models share the fundamental idea with this framework that such *good* representations must be learnt via prediction in a flexible, learnable latent space. In this context, the same predictive principle should be generally applied throughout the whole speech cortex hierarchy, from the lowest to higher-levels speech representations. In this regard, the wav2vec 2.0 features were high-dimensional speech signals, possibly noisy and redundant in the context of a temporal prediction task, that can be considered intermediate representations within the broader hierarchical processing stream^21^. Our findings show that, when treating these features as fixed, optimizing predictive information through prediction error minimization of future inputs does not yield behaviorally relevant representations. Instead, our results reinforce the idea that the brain extracts predictive information through prediction in a learnable latent space, where potentially irrelevant, high-variance details can be filtered out. This directly links to current problems in machine representation learning^57^ which share a similar goal with our nervous system: building representations that contain information readily useful for further downstream tasks. Indeed, in this field, successful learning depends not only on model size and training data but, critically, on how relevant information is represented, an aspect determined by the learning objective itself. Two main classes of approaches have emerged in machine learning: generative architectures and joint-embedding architectures (JEAs)^58^. Generative models, such as our predictive autoencoder (deep-pAE), learn by reconstructing observed inputs, whereas JEAs learn representations directly in latent space without requiring accurate reconstruction. Our contrastive learning (CL) model belongs to this type of learning algorithms, specifically trained with an objective function that optimizes the selection of predictive information (InfoNCE)^*59*^. In temporal prediction settings, JEAs training involves three steps: (i) encoding observations into abstract latent representations, (ii) predicting future latent states, and (iii) evaluating predictions against actual future representations. Interestingly, this procedure is reminiscent of Craik’s three-steps algorithm for internal model fitting.^1^

This parallel offers a way for the investigation of the computational principles underlying neural processing of sensory signals. In conclusion, by explicitly comparing different efficient information coding hypotheses, our results indicate that representations optimized for predictive information via prediction in latent space better account for both neural activity and perceptual behavior during speech processing. More broadly, we suggest that framing neural representation learning as a problem of objective function optimization provides a powerful and controlled approach to testing normative theories of perception. Future models that more directly align their learning objectives with biological constraints and goals, particularly efficient representation of predictive information, explicit representation of redundant and non-redundant information in the signals, may not only improve machine representation learning performance but also offer deeper insight into how meaningful perceptual representations emerge from natural sensory signals.

## Materials and Methods

### Dataset

#### EEG and behavioral data

We recorded and analyzed EEG activity from 26 healthy Italian participants while they listened to English sentences. EEG data were continuously recorded using a 64-channel active electrode system (BrainAmp MR Plus; Brain Products GmbH, Gilching, Germany). Each participant listened to 100 sentences from the USC-TIMIT database^29^. Each sentence was presented three times, resulting in around 17 minutes of active listening per participant. A rhyming task followed each sentence. Additionally, participants completed a behavioral session that included a reading and a transcription task. In the reading task, each participant read aloud 50 of the 100 selected sentences.

In the transcription task, participants listened to the remaining 50 sentences and transcribed them to the best of their ability. This design ensured that each participant either read or transcribed each of the 100 sentences, resulting in 50 transcripts and 50 audio recordings per participant.

#### Stimuli

Stimuli were selected from the USC-TIMIT dataset, which includes synchronized recordings of audio and articulatory movements from 10 native American English speakers, along with transcripts. Audio was recorded at a sampling rate of 22.05 kHz, and articulator kinematics (lips, jaw, tongue) were tracked at 100 Hz using an electromagnetic articulography (EMA) system: the NDI Wave Speech Research System (Northern Digital Inc., Waterloo, Ontario, Canada). We used 100 sentences spoken by the same male speaker (M1 in the dataset), with durations ranging from 2.9 to 4.5 seconds. Silent segments at the start and end were removed, and all audio stimuli were normalized to the same average intensity (rms = 0.0316). Transcripts and audio stimuli were presented to participants during the experiment.

#### Rhyming task

For each sentence, four words (two rhyming and two non-rhyming) were selected. Rhyming words were identified using an online rhyme generator (https://www.rhymer.com/), and non-rhyming words were selected using a random word generator (https://ohmyluck.com/it/random-word/). To prevent response bias, rhyming and non-rhyming words were matched for frequency in English using the wordfreq Python library (https://pypi.org/project/wordfreq/) resulting in an average frequency of ~6e-05. In each trial, one word was randomly chosen from the four-word list and displayed, ensuring no word repetitions.

### Experiment

#### Participants

Thirty-one healthy, right-handed Italian native speakers with normal or corrected-to-normal vision participated in the study and were compensated 40 euros. All participants self-reported English as their second language. Written informed consent was obtained, and the study was approved by the local ethical committee, “Comitato Etico Unico della Provincia di Ferrara” (approval no. 170592). One participant was excluded due to technical issues, and four more were excluded during preprocessing, leaving 26 participants (16 females; age: 21.96 ± 1.95 years).

#### Behavioral setup and procedure

The behavioral session occurred two days before the EEG session and consisted of a reading and transcription task. Participants sat 60 cm from an LCD monitor with two loudspeakers positioned 20 cm from each side. The session lasted approximately 45 minutes, including instructions.

Reading Task: A microphone recorded participants reading 50 randomly selected sentences out of the 100 stimuli at a sampling rate of 44.1 KHz. An algorithm implemented in the MATLAB environment detected speech onset and offset to manage recording start and end. After a brief pause, the next sentence appeared written on screen.

Transcription Task: Participants listened to the remaining 50 sentences and typed what they heard, with a 30-second timeout. The transcription task always followed the reading task to avoid bias in the manner of production.

#### EEG setup and procedure

EEG data were recorded continuously during the experiment with a 64-channel active electrode system (BrainAmp MR Plus; Brain Products GmbH, Gilching, Germany). Electrooculograms (EOGs) were recorded with four electrodes from the cap (FT9, FT10, PO9 and PO10) that were removed from the original scalp sites and placed bilaterally at the outer canthi and below and above the right eye to record horizontal and vertical eye movements, respectively. All the electrodes were online referenced to the left mastoid. The impedance of the electrodes was kept below 15 kOhm. EEG signals were acquired at 1,000 Hz. During the experiment, participants sat in front of an LCD monitor (VIEWPixx/EEG; 24 in., 120 Hz) at approximately 80 cm and were instructed to put their hands on two pads placed on their right and left sides. Two loudspeakers were placed at 20 cm from each side of the screen.

The session consisted of 300 trials divided into three blocks of 100 trials and split into two sub-blocks of 50 trials each, with short in-between breaks. In each block, all the stimuli dataset was presented in a random order, so that every stimulus was heard three-times by each participant at the end of the experiment. In each trial, the following sequence of events occurred: 1) participants were presented with a black fixation cross at the center of a uniformly colored gray screen; 2) after a variable time (between 1 and 2 s), the audio of a randomly selected sentence was played; 3) after a variable time (between 1 and 2 s) from the end of the acoustic stimulus, a word appeared at the center of the screen in place of the fixation cross; 4) participants had to indicate whether or not the word presented rhymed with one of the words in the previously heard sentence by pressing one of the two pads; and 5) the trial ended when the participants gave a response or after a time-out of 10 s if no response was given. Participants were asked to reduce their blinks as much as possible and to maintain their eyes on the fixation cross for the entire duration of the sentence.

The entire experiment lasted ~2 h, including the EEG cap mounting and preparation. Stimulus presentation and button-press acquisition were controlled via MATLAB (The MathWorks, Inc.; https://www.mathworks.com; RRID:SCR_001622) and the PsychToolbox-3 extensions (http://psychtoolbox.org; RRID:SCR_002881). All relevant events in the trial (i.e., trial start, stimulus onset, stimulus offset, word appearance, and button press) were converted into a TTL (transistor-transistor logic) by the VIEWPixx/EEG system to accurately synchronize them with the EEG data. The audio tracks were simultaneously sent to the loudspeakers for stimulus presentation and to the EEG device using an analog port to ensure the alignment between neural and acoustic recording.

## Data Analysis

Analyzes were performed within the Python computing environments using open-source libraries: MNE (https://mne.tools/stable/indexhttps://mne.tools/stable/index.html; RRID:SCR_005972), GCMI (https://github.com/robince/gcmi), CEBRA (https://github.com/AdaptiveMotorControlLab/CEBRA), as well as custom made code (https://github.com/ctnsc-unife/JEACL-neural-principles). We, furthermore, used the open-source model wav2vec 2.0 as available in the torchaudio bundles (https://pytorch.org/audio/main/generated/torchaudio.models.Wav2Vec2Model.html) and the Open AI’s text embedding model text-embedding-ada-002 (https://platform.openai.com/docs/guides/embeddings). We will firstly approach the presentation and motivation of the general framework of the analyses and then go further with the precise methodological description of the analysis pipeline.

### General framework

When processing speech stimuli, the human brain transforms the observable auditory input into multiple latent representations that facilitate speech comprehension. The exact transformations applied by the brain are largely unknown but are reflected in the recorded EEG signals during the conducted experiment. We, then, model speech processing to extract latent representations using a fixed artificial neural network (ANN) backbone optimized for speech recognition, namely the wav2vec 2.0, and a learnable joint-embedding architecture that uses contrastive learning (CL), or deep autoencoders (deep-AE and deep-pAE) on top of it. We aimed to compare latent representations extracted by the models with those extracted by the participants’ brains listening to the same sentences. To do so, we compute mutual information (MI) between latents inferred by the models and the participants’ EEG signals. The higher the MI values, the more model-extracted features align to the brain’s latent representations, reflecting stronger correspondence between the modeled and the real neural processes underlying speech perception.

### Behavioral data analysis

The speech transcription task from the behavioral session was analyzed to assess participants’ speech recognition performance in their second language (English). During the task, each participant transcribed 50 random sentences out of the 100 selected acoustic stimuli. To quantify the accuracy of these transcriptions, both the participant-written sentences and the correct transcripts were encoded using OpenAI’s text embedding model, text-embedding-ada-002. This pre-trained model generates dense, multidimensional vector representations that capture the semantic meaning of text, making it particularly suitable for similarity analysis.

Each written sentence was converted into a 1,536-dimensional vector in this high-dimensional space, where correctly identified transcripts were encoded similarly to the original ones. In this space, vectors closer together indicate greater semantic similarity, while those farther apart indicate greater dissimilarity. As recommended in the model documentation (https://platform.openai.com/docs/guides/embeddings), cosine similarity was used to calculate the similarity between each participant’s transcription and the corresponding correct transcript. To evaluate each participant’s overall transcription performance, the average similarity across all 50 sentences was calculated. This score reflects the accuracy of each participant’s transcriptions, with higher values indicating better performance.

### EEG preprocessing

Continuous EEG data were preprocessed following standard procedures. The signals were band-pass filtered between 0.2 and 45 Hz using a two-pass, 4th-order Butterworth filter. The data were then down-sampled to 100 Hz and re-referenced to the common average. Speech onset was detected using the acoustic track recorded alongside the EEG data. A custom Python code identified the speech onset times, which were used to epoch the EEG data. Epochs were aligned to the acoustic stimulus, spanning from 1 second before to 5.5 seconds after stimulus onset, which is 1 second after the end of the longest sentence. Noisy trials were identified through visual inspection and removed. Artifacts caused by eye movements and heartbeat were addressed using Independent Component Analysis (ICA) with the fastica algorithm as implemented in the MNE library. Noisy channels were excluded from the ICA computation and subsequently replaced by linear interpolation of neighboring channels. After preprocessing, a total of 288.5 ± 13.5 trials (mean ± SD) were retained for further analysis.

### Speech encoding model: wav2vec 2.0 features

Speech features were extracted using wav2vec 2.0, a state-of-the-art pre-trained model for speech technologies^19^. While originally designed for speech recognition tasks, we employed wav2vec 2.0 exclusively as a feature extractor to convert raw audio waveforms into multidimensional representations of speech. The wav2vec 2.0 architecture consists of two primary components:

Feature Encoder: A multi-layer convolutional module that processes audio input to generate latent representations. Due to the convolutional operations, the audio signal is effectively resampled to approximately 50 Hz.

Transformer Encoder: A self-attention-based module that integrates information across the entire speech sequence, producing 12 context representations, each with 768 dimensions.

For this study, we used the WAV2VEC2_ASR_BASE_960H model available in the torchaudio pipelines package (https://pytorch.org/audio/main/generated/torchaudio.models.Wav2Vec2Model.html). This model was pre-trained on 960 hours of unlabeled audio from the LibriSpeech dataset and fine-tuned on the corresponding transcripts. To prepare the 100 speech stimuli from the USC-TIMIT dataset for feature extraction, each stimulus was resampled to match the model’s required input sampling rate of 16 kHz. The stimuli were then passed through the wav2vec 2.0 model, and only the final (12th) context representation was extracted for further analysis.

### Speech encoding model: Dimensionality reduction with PCA

To reduce the dimensionality of wav2vec 2.0 features and approximate the process of extracting relevant information, we employed Principal Component Analysis (PCA), a widely used dimensionality reduction method in neuroscience. PCA operates on the principle that relevant information is proportional to the variance in the input data. Since large portions of variance in a dataset are often captured within a small subset of dimensions, PCA effectively reduces dimensionality while retaining most of the essential information. PCA was applied to the matrix D of concatenated wav2vec 2.0 speech representations. PCA works by performing eigendecomposition of the covariance matrix Σ_D_, identifying a new coordinate system represented by eigenvectors. These eigenvectors correspond to the principal components (PCs), with each component ordered by the amount of variance it captures. The eigenvalues of Σ_D_ quantify the variance explained by each principal component, making it possible to prioritize dimensions that capture the most variance.

Dimensionality reduction using PCA was achieved using targeted cumulative explained variance ratios:

1. Top subspaces: Retaining components cumulatively explaining 25%, 50%, 75%, or 99% of the variance starting from the first PC.
2. Bottom subspaces: Retaining components cumulatively explaining 0.1%, 1%, 25%, or 50% of the variance, starting from the lowest-variance components.

For both strategies, the explained variance ratio was calculated by summing the contributions of the eigenvalues in the desired order (highest or lowest variance) to determine the number of PCs required to achieve the targeted variance ratio.

### Speech encoding model: Contrastive learning in a joint-embedding architecture

We used the context features from the wav2vec 2.0 model as representations of speech signals. These features are high-dimensional (768 × time length) and may contain significant redundancy. In contrast, the brain processes sensory inputs by extracting relevant information and compressing it into efficient, low-dimensional neural representations. To model this property, we employed contrastive learning (CL) within the framework of joint-embedding architectures (JEA). This approach helps to filter out irrelevant information by capturing shared information between consecutive speech samples while optimizing the InfoNCE loss function, which, in the limited sample case is defined as:

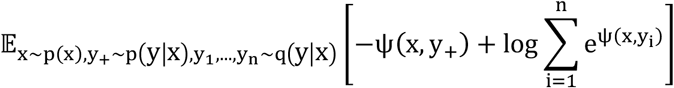

Here, *x* is the reference sample, y_+_ is the positive sample and y_1_, …, y_n_ are the negative samples. The function ψ(·) measures similarity and is defined as:

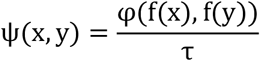

where *τ* is the temperature parameter, φ is a similarity measure (e.g., cosine similarity), and f(·), is a parametric function. Each stimulus was represented as a matrix of shape (768, Nsamples_i_) where Nsamples_i_ corresponds to the number of time samples in the i-th stimulus, recorded at 50 Hz (the wav2vec 2.0 output sample rate). We concatenated all stimulus matrices across the time dimension into a single matrix D of shape (768, 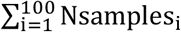). This matrix served as the input dataset for training the CL model within the CEBRA framework^28^. To train the model:

1. A reference sample x of shape (768,) was randomly selected, and its immediate subsequent sample was designated as the positive sample y_+_. 4096 negative samples y_1_, …, y_4096_ were randomly drawn from the dataset, thus using a uniform prior.
2. Both *x, y*_+_, and *y*_1_, …, *y*_4096_ were processed by a shared encoder, a multi-layer perceptron (MLP) referred to as the “offset1-model-v3” in the CEBRA framework. This encoder consists of three linear layers, each with 128 hidden units and GELU activations. Skip connections are implemented to improve training stability.
3. The encoder implements the parametric function *f*(·) and projects the data into a normalized K-dimensional latent space using a final linear layer.
4. The InfoNCE loss was calculated using cosine similarity as the metric *φ* and a fixed temperature parameter *τ* = 1, in batches of 4096 references-positives pairs.
5. The model was trained for 10000 steps with a batch size of 4096 and a learning rate of 0.0001 using the Adam optimizer. Training was conducted on a CUDA-enabled GPU for efficiency.

The following code snippet initializes one model and training parameters in the CEBRA framework:

~~~
import cebra
from cebra import CEBRA
# Initialize the CEBRA model
CL_model = CEBRA (
  model_architecture=‘offset1-model-v3’,
  batch_size=4096,
  temperature=1,
  learning_rate=1e-4,
  max_iterations=10000,
  time_offsets=1,
  # 20ms offset optimizer=“adam”,
  output_dimension=7,
  num_hidden_units=128,
  device=“cuda”,
  conditional=‘time’,
)
~~~

To model brain processes operating at different temporal scales, we trained 10 distinct models, each with a progressively larger time_offsets parameter. The parameter ranged from 1 to 10, corresponding to positive samples taken between 20 ms and 200 ms ahead of the reference sample. This allowed us to investigate the impact of varying temporal predictions on the learned representations.

### Speech encoding model: Goodness-of-fit in contrastive learning models

In the CEBRA framework, the InfoNCE loss can be used as a goodness-of-fit metric to compare models at the same point in training time. Indeed, the infinite samples loss converges to the expression:

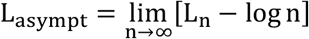

Where L_n_ is the approximated limited sample loss and n is the batch size. This convergence is constrained by the following inequality^28^:

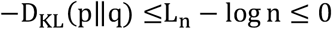

here, D_KL_(p‖q) represents the Kullback-Leibler divergence between the distributions p and q. This asymptotic loss can serve as a measure of model goodness-of-fit when compared under identical training conditions (i.e., same number of steps). A lower value of L_n_ − log n indicates better model fitting^28^. To ensure the goodness-of-fit metric is always positive, we used instead the opposite value log n − L_n_, where higher values indicate better fitting.

This metric was employed to select the optimal latent dimensionality for the model. We trained 18 different CL models, each differing in the dimensionality of the latent space. The dimensions tested were: 2,3,4,5,6,7,8,9,10,11,12,13,14,15,16,32,64, and 128.

At the end of training, we computed the average log n − L_n_ over the last 100 training steps and employed the measure as goodness-of-fit for each model. This allowed us to systematically evaluate the effect of latent dimensionality on the model’s performance and select the dimensionality that provided the best balance between fitting and compression. We therefore did not use this metric to test hypotheses in a supervised setting as suggested by Schneider and colleagues^28^, but rather to minimize the number of output dimensions of the model and produce low-dimensional latents.

Apart from this optimal dimensionality, we also trained models at other various output dimensionalities: 3, 16, 32, 64, 128.

### Speech encoding model: InfoNCE in CL as a measure of predictive information

As originally proposed by van den Oord and colleagues^60^, the InfoNCE loss can function as a mutual information (MI) estimator. Specifically, given x and y_+_ as the reference and positive samples, respectively, and their corresponding latent representations u = f(x) and v = f(y_+_), the following relationships hold^60^:

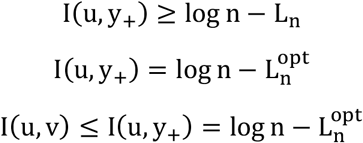

Here, I(u, y_+_) denotes the mutual information between the latent representation of the reference sample u and positive sample y_+_. The metric log n − L_n_ serves as a lower bound on this mutual information, while at the minimizer of the InfoNCE loss 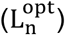, it becomes an upper bound on the mutual information between u and v (the latent representations of the reference and positive samples). We leveraged this property to estimate the amount of information that the model can extract from the reference speech sample (x) that is relevant for predicting the positive speech sample (y_+_). This estimate provides insights into the model’s ability to encode meaningful and predictive features from the sensory input.

Even if InfoNCE loss does not incorporate an explicit regularization term to constrain the amount of information retained in the latent space, it has been shown that it overfits to the shared information between reference and positive sample, thus approximately retrieving a minimal sufficient statistics^61^. A tentative explanation of the phenomenon is given in section 4.1 of Schwatz-Ziv and LeCun^63^.

### Speech encoding model: Classifying samples based on predictive information

Building on the use of InfoNCE as a mutual information estimator, we employed the value of the InfoNCE loss after training to assess the model’s ability to extract predictive information from a reference sample about a corresponding positive sample.

To estimate the InfoNCE loss reliably, we sampled one reference, one positive, and 4096 negative samples, as done during training. To quantify the information extracted by the model about each speech sample at varying time lags, we used the matrix of concatenated stimuli D. The positive sample was fixed, while the reference sample was shifted backward (negative lags) and forward (positive lags) in time: I(u_t+k_, y_t_) for k=−0.6, −0.58, …, 0.8 seconds, with 20 ms steps. The model trained with 20 ms time_offsets was used for these computations. This process was repeated across all samples in D, and the results were averaged over all positive samples. The mutual information measure was expected to peak at lag 0, where the reference and positive samples coincide. To investigate the predictive information accumulated about a positive sample within the preceding 200 ms time window, we defined a ΔI metric as follows:

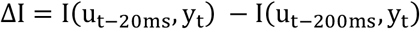

The ΔI value was computed using the InfoNCE loss during inference as described. We then classified positive samples (y_t_) into two groups based on their ΔI values:

1. Low-Δ information samples (LΔI): Corresponding to the first quartile of the ΔI distribution.
2. High-Δ information samples (HΔI): Corresponding to the last quartile of the ΔI distribution.

LΔI samples accumulate minimal information during the 200 ms preceding their presentation, while HΔI samples accumulate substantial information within this time frame. Despite this difference, both classes become almost equally predictable at the time of their presentation (k=0; see below).

### Speech encoding model: Comparing predictive information patterns of LΔI and HΔI classes

We computed mutual information separately for the LΔI and HΔI classes over time:

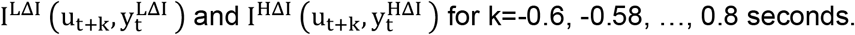

Here, 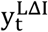 and 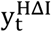 represent samples belonging to the LΔI and HΔI classes, which do not overlap. Given their classification, LΔI samples were more predictable than HΔI samples at lag of −200 ms.

We then computed the average δ-information for the two classes by evaluating the information difference between consecutive time steps:

For LΔI:

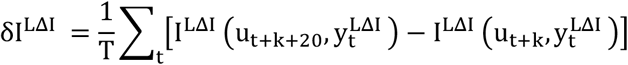

For HΔI:

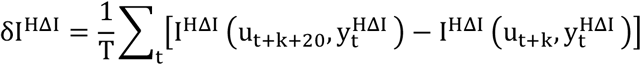

These measures allowed us to compare how predictive information was accumulated for both low-Δ information and high-Δ information samples over time in the listeners’ brain.

### Speech encoding model: Reconstruction-based deep-autoencoders

As done for contrastive learning models, we also trained reconstruction-based deep-autoencoders on the same dimensionality reduction task. Specifically we trained two different types of deep-autoencoders: i) an autoencoder that takes as input only reference samples at every training step (deep-AE), it projects the samples into a low-dimensional latent space, and then reconstructs the original input samples through a decoder network; ii) an autoencoder that takes as input both reference and positive samples at every step (deep-pAE), it encodes references into a compressed latent space, and implements a decoder to reconstruct the positive samples.

As with the CL models and PCA, the deep-AE and deep-pAE were trained on the matrix D of concatenated wav2vec 2.0 speech features. The architecture of the proposed autoencoder leveraged the encoder design for the CL, comprising:

- A three-layer perceptron with 128 hidden units per layer.
- GELU nonlinearities between layers to introduce nonlinearity.
- Skip connections to facilitate efficient training.

To ensure comparability with the CL, we implemented a symmetric decoder tasked with reconstructing the speech wav2vec 2.0 signals at the subsequent time step relative to the input. The autoencoder was trained for 10,000 steps with the following parameters:

- Batch size: 4096 samples.
- Learning rate: 0.0001
- Loss function: Mean Squared Error (MSE).
- Optimizer: Adam.

For the deep-pAE training, positive samples were sampled using the same conditional distribution used for CL training, namely using a fixed time offset, which in our case equals 1.

By employing the same encoder structure as the CL and targeting reconstruction of the subsequent time step, we maximized the similarity between the deep-pAE and CL models training. This setup allowed a robust comparison of dimensionality reduction and relevant information extraction methods.

### Decoding analyses

We evaluated models’ skills in three interesting linear decoding tasks: i) sample prediction in latent space, ii) sample reconstruction in observed space, and iii) sample prediction in observed space. These are the task for which CL, deep-AE, and deep-pAE are optimized during training, respectively. To do so we trained linear regressions using a 5-fold validation scheme using a test set of 20% of dataset at every fold and computed average R^2^ values. Specifically, given speech reference x, positive y_+_, latent representation of reference *u* and of positive *v*, the latent space prediction task entails fitting a linear model to predict *v* from *u*; the reconstruction task requires linear prediction of x from *u*; while the prediction in observed space means predicting y_+_ from *u*. these analyses are separately repeated for models of various output dimensionalities: 3, 7, 16, 32, 64, 128.

We further tested the models at a sentence ID decoding task. In this case we employed the same 5-fold cross-validation scheme while using a kNN decoder to accomplish the task. In this case, an additional validation set is used to tune the neighbors parameter k of the decoder. Specifically, speech samples were randomly drawn from the output of each of the three models (CL, deep-AE, deep-pAE) together with the ID of the sentence they were coming from. The kNN decoder is trained to predict the ID from the speech outputs. Since there are 100 sentences in the dataset, the chance in this task is 1% decoding accuracy.

### EEG encoding of latent features

To assess the brain-relevance of the extracted latent speech representations (PCA, CL, CL-LΔI, CL-HΔI, deep-AE, deep-pAE), we quantified the alignment between these representations and the EEG signals using mutual information (MI)^64^. A higher MI indicates stronger alignment, suggesting that the latent features capture information akin to that processed by the brain during speech listening. MI measures the reduction in uncertainty about a random variable X given knowledge of another variable Y. It serves as a statistical test against the null hypothesis that X and Y are independent, capturing both linear and nonlinear relationships. In simpler terms, MI quantifies the degree to which the variability of X can be predicted by observing Y. We employed the Gaussian Copula Mutual Information (GCMI) estimator to calculate MI. GCMI provides a robust lower bound on the true MI value and is particularly suited for datasets of limited size^31^. The GCMI method involves the following steps:

1. The EEG recordings were resampled to 50 Hz to match the sampling rate of the latent speech representations.
2. Both EEG signals and latent features were band-pass filtered between 0.5 and 24 Hz using a second-order Butterworth filter (two-pass to avoid phase shifts).
3. EEG signals were shifted in time relative to the speech representations, ranging from −0.6 to 0.8 seconds, where negative lags represent anticipatory brain activity, while positive lags correspond to activity following stimulus listening.
4. Both variables were transformed through copula normalization, which preserves the statistical dependency (copula) between variables while normalizing their marginal distributions to be univariate Gaussian. Mutual information remains unchanged after this transformation, as it depends solely on the copula linking *X* and *Y*. The mutual information *I*(*X, Y*) can be expressed in terms of entropy as:

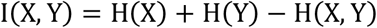

Where H denotes entropy. Using the concept of copulas, this becomes:

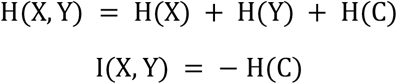
5. Finally, MI was computed using the analytical expression for Gaussian variables:

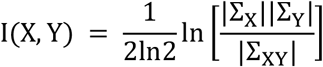

Where Σ_X_ and Σ_Y_ are the covariance matrices of X and Y and Σ_XY_ is the covariance matrix of their joint variable. This calculation was performed for each EEG channel and group of latent representations using the GCMI function (gcmi_cc).

### Statistical Analyses

To determine whether the speech latents (wav2vec 2.0 CL, PCA, deep-AE) were significantly encoded in the EEG recordings, the mutual information (MI) outputs were evaluated against surrogate data via circular shift. The surrogate data disrupted the original relationship between the EEG signals and the speech latents while preserving the statistical properties of each signal, including its autocorrelation^65^. Specifically, the EEG activity at each electrode and trial (epoched and bandpass filtered as described previously) was circularly shifted by a random number of samples, selected between N/3 and N – N/4, where N represents the number of samples in the shortest trial (296 samples). As in the original analysis, MI was computed between the EEG and each group of latents. This procedure was repeated 100 times to generate a surrogate distribution for each participant and each information component. One-tailed cluster-based permutation statistics^32^ were then applied to test at the group level whether the original MI values were significantly greater than the mean of the surrogate distribution. The null hypothesis assumed that the real and surrogate MI values belonged to the same probability distribution. A nonparametric statistical test was used to avoid assumptions about the distributions of the information values. All MI output samples (one per condition for each participant) were combined into a single set and randomly partitioned into two subsets. The test statistic was calculated by employing a univariate statistical test to compute a lag-specific t-value (using the formula for dependent samples). Lags with absolute t-values exceeding a predefined threshold were grouped into clusters based on temporal adjacency. The threshold for clustering was set to the 95th quantile of the T-distribution, corresponding to a critical alpha level of 0.05. For each cluster, the cluster-level statistic was calculated by summing the t-values of all lags within the cluster. This procedure was repeated 5000 times with random permutations to construct a distribution of the largest (absolute) cluster-level statistics. The test statistic was then computed for the non-permuted data (real and surrogate) to identify clusters of lags, and their p-values were determined as the proportion of random permutations with a larger (absolute) test statistic than the non-permuted data (Monte Carlo estimate). This method inherently controls the false alarm (FA) rate, setting the probability of Type I errors to the critical alpha level (0.05), and addresses the multiple comparison problem (MCP) by performing hypothesis testing at the cluster level. To compare the MI values of CL-LΔI to CL-HΔI, as well as comparing CL vs deep-AE vs deep-pAE encodings, the same nonparametric statistical test was applied. Surrogate data were not required for this comparison, as the distributions of the two classes were directly compared using a two-tailed cluster-based permutation test. In the case of CL-LΔI and CL-HΔI, we specifically considered constraints on the model’s δ-information encoding preceding stimulus presentation (anticipatory activity). To account for a biological lag in information processing, we cross-correlated the δ-information pattern in both classes with their EEG information processing. For the statistical analysis, only channel-averaged MI values corresponding to valid lags (≤ peak) were included.

## Extended Data

**Figure 1.**
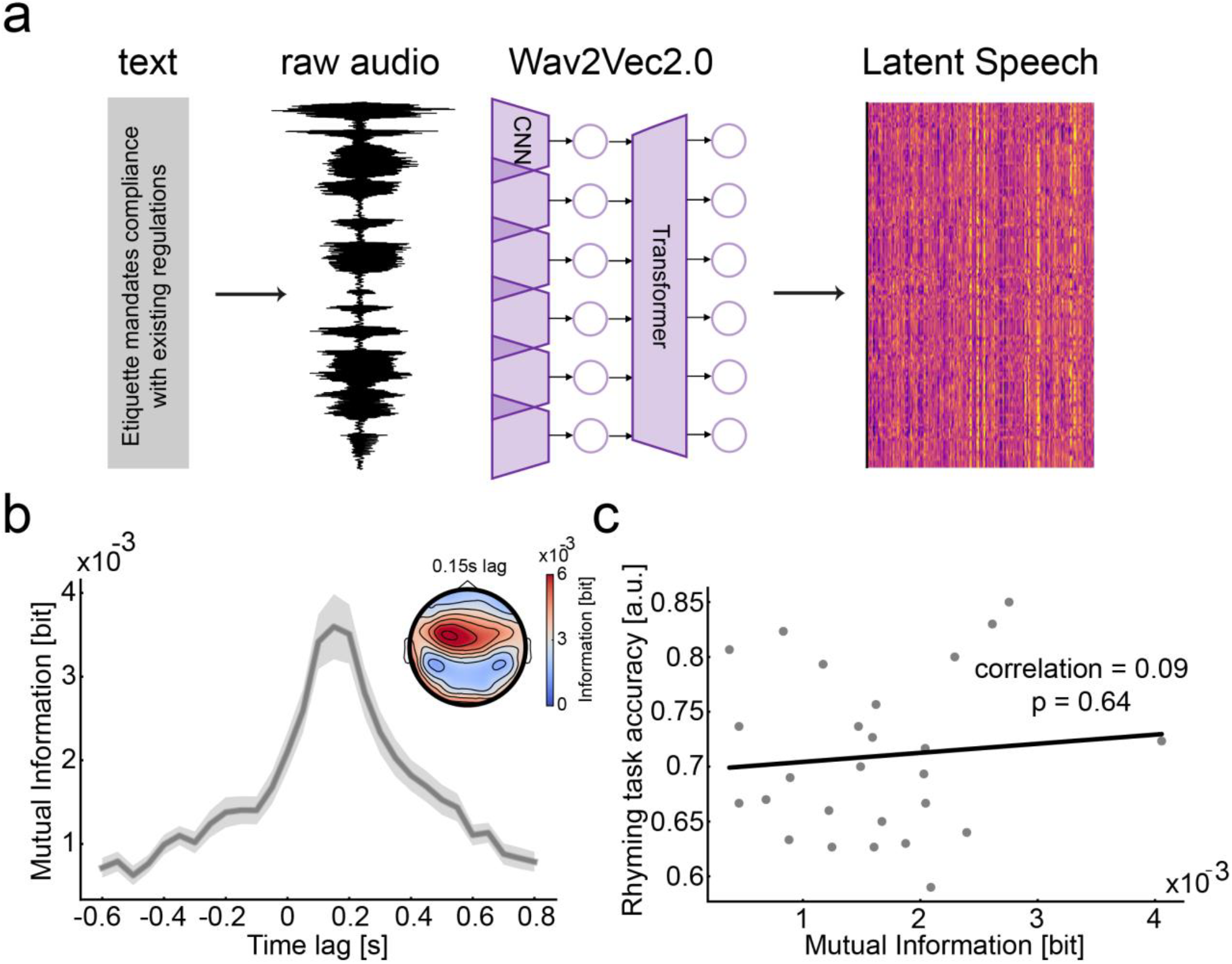
wav2vec 2.0 latent features do not explain participants’ speech recognition performance. **a**. Schematic of the extraction of the wav2vec 2.0 features. Textual information is linked to acoustic recordings which are transformed by wav2vec 2.0 base model to produce 768-dimensional latent speech features useful to accomplish speech-to-text tasks. **b**. Mutual information (MI) computed between wav2vec 2.0 latent speech features and EEG activity of participants listening to the same sentences. Latents information is significantly encoded across all the tested lags (p = 2e-04; one tail cluster-based statistics against the mean of 100 surrogate data). **c**. Scatter plot of MI values against rhyming task accuracy for each participant. No significant correlation is found (*r* = 0.09; *p* = 6.4e-01).

**Figure 2.**
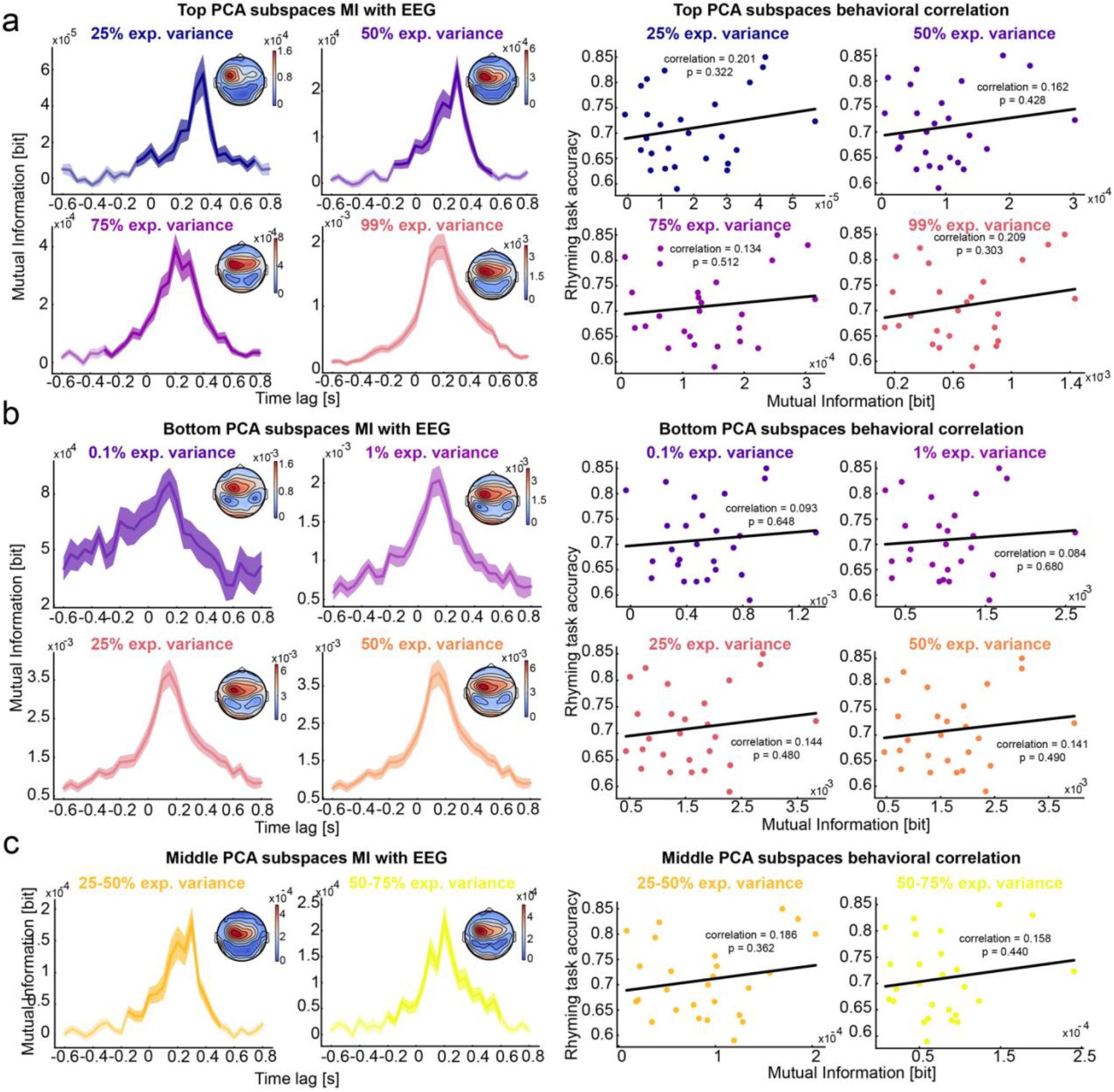
EEG encoding of wav2vec 2.0 PCA subspaces and correlation with rhyming task accuracy. **a**. Top PCA subspaces analysis. Top PCA subspaces are computed by accumulating explained variance ratio starting from the first PC (PC1). Though information is significantly encoded in the EEG activity at all tested subspaces, no correlation is found with participants’ rhyming task performance. **b**. Bottom PCA subspaces analysis. Bottom subspaces are computed by accumulating explained variance ratio starting from the last PC (PC768). Also in this case, EEG encodes information about these subspaces, but no significant correlation is found. **c**. The same analyses are repeated for some middle explained variance PCA subspaces and the same results are found.

**Figure 3.**
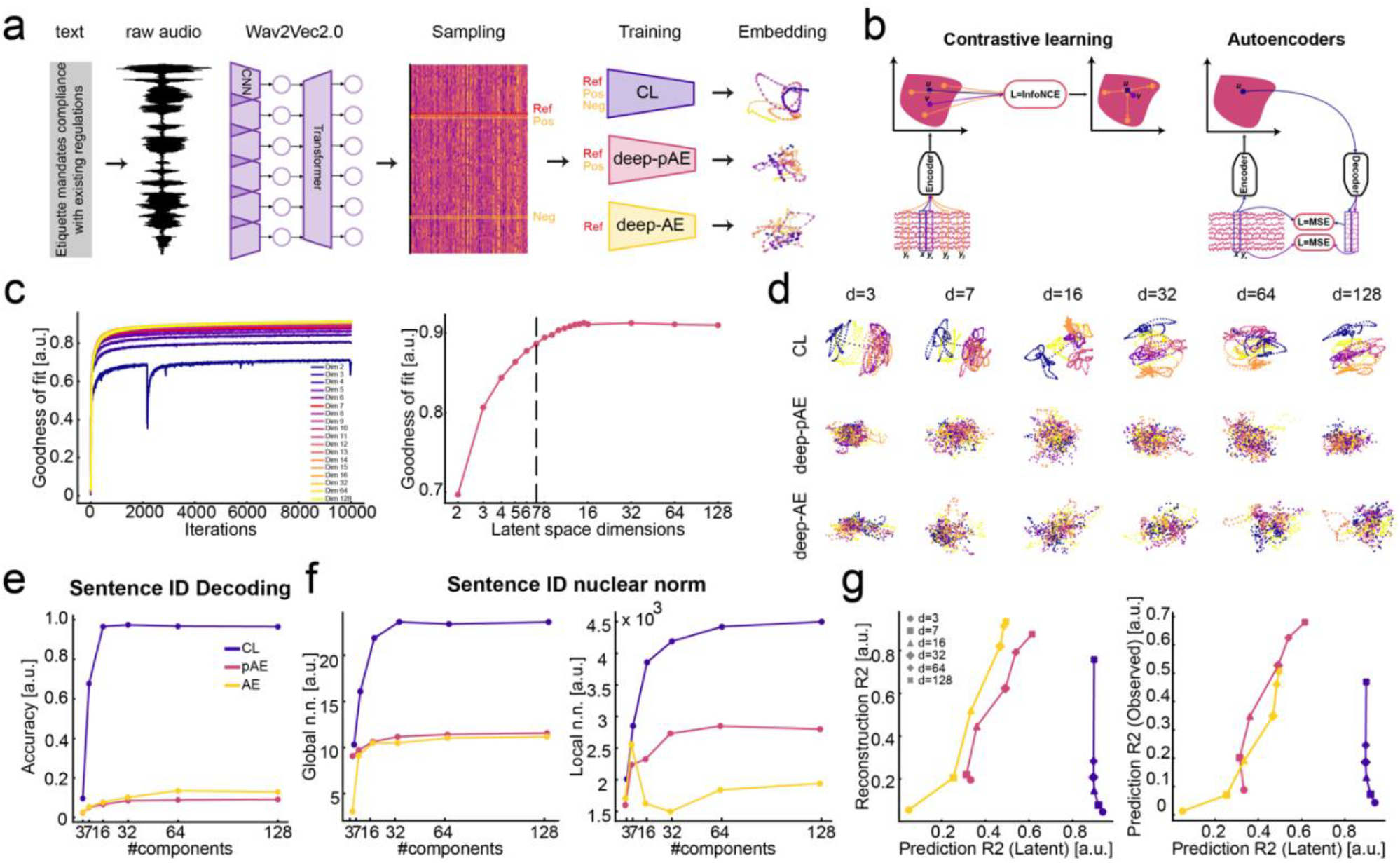
Nonlinear dimensionality reduction of wav2vec 2.0 features. **a**. Dimensionality-reduction pipeline: (i) each sentence is paired with its raw audio; (ii) audio is encoded by pretrained wav2vec 2.0 into 768-dimensional features; (iii) these features train the three architectures (CL uses reference + positive + negative samples; deep-pAE uses reference + positive; deep-AE uses reference only); (iv) trained models produce latent embeddings for each sentence. ***b***. *Precise s*chematic of the two alternative approaches used to solve dimensionality reduction for relevant information extraction. **c**. Selection of output dimensionality based on goodness-of-fit. **d**. Example latent spaces for five sentences. Only CL maintains clear sentence clustering even at small dimensionalities. **e**. Sentence-ID decoding using a kNN classifier (chance = 1%). Despite no task-specific supervision, CL far outperforms both generative models. **f**. Global and local nuclear norms treating sentences as classes. CL exhibits higher global separability and richer (less compressed) local structure. **g**. Signal reconstruction vs. prediction. deep-AE excels at reconstruction but fails at prediction; deep-pAE best predicts future signals; CL uniquely optimizes latent-space prediction, improving reconstruction only at higher dimensionalities.

**Figure 4.**
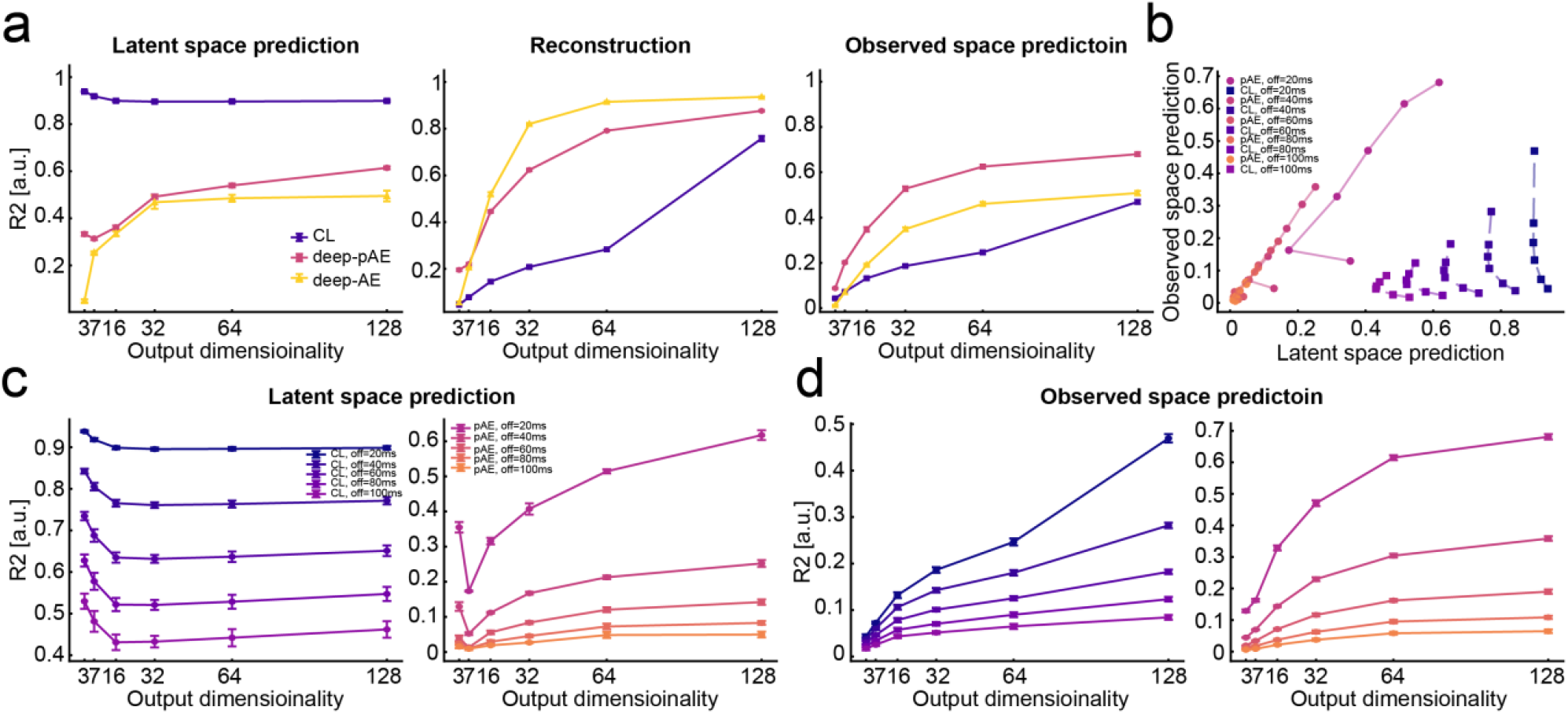
ANN modeling enables testing different speech perception hypotheses. **a**. R^2^ values of the three models (CL, deep-AE, deep-pAE) at three tasks: latent space prediction, reconstruction, observed space prediction. CL excels at latent space prediction, deep-AE is the best at reconstruction, deep-pAE is the best at observed space prediction. **b**. R^2^ latent space prediction values for CL and deep-pAE across various temporal offsets (from 20 to 100 ms ahead in time). **c**. R^2^ observed space prediction values for CL and deep-pAE across various temporal offsets (from 20 to 100 ms ahead in time). **d**. Comparison of the two models in the two prediction tasks. CL remains consistently good at solving latent space prediction even for longer temporal offsets, while deep-pAE drastically drops its observed space prediction performance as soon as the temporal offset is increased.

**Figure 5.**
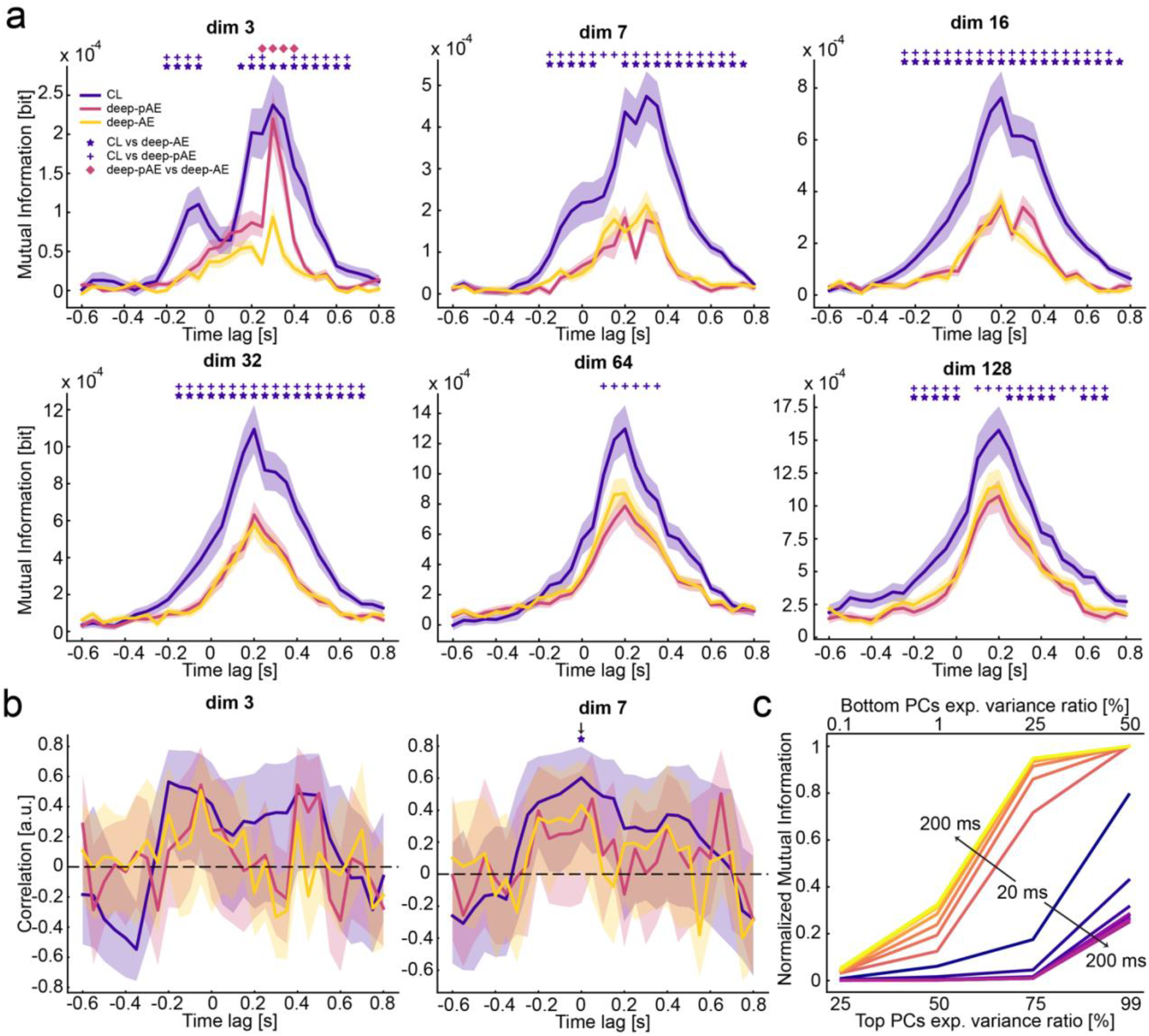
Contrastive learning features are more encoded in EEG activity and correlated with rhyming task accuracy. **a**. MI values between EEG activity and latent features extracted from CL, deep-AE, and deep-pAE computed in a time-resolved fashion from −0.6 to 0.8 s lag. Results are computed and plotted for all tested model output dimensionalities. CL latents are consistently better encoded than the other two output latents. CL vs p-AE (dim 3: from −0.2s to −0.05s *p* = 1.11e-02 and from 0.2s to 0.25s *p* = 3.98e-02 and from 0.4s to 0.65s *p* = 1.20e-03, dim 7: from −0.15s to 0.7 *p* = 2.00e-04, dim 16: from −0.25s to 0.7s *p* = 2.00e-04, dim 32: from −0.25s to 0.75s *p* = 4.00e-04, dim 64: from 0.1s to 0.35s *p* = 2.22e-02, dim 128: from −0.2s to 0s *p* = 2.40e-02 and from 0.1s to 0.7s *p* = 3.40e-03; two-tailed cluster-based statistics) and CL vs AE (dim 3: from −0.2s to −0.05s *p* = 5.6e-03 and from 0.15s to 0.65s *p* = 2.00e-04, dim 7: from −0.15s to 0.05s *p* = 2.1e-02 and from 0.2s to 0.75s *p* = 2.00e-04, dim 16: from −0.25s to 0.75s *p* = 2.00e-04, dim 32: from −0.15s to 0.75s *p* = 2.00e-04, dim 128: from −0.2s to 0s *p* = 4.24e-02 and from 0.25s to 0.45 *p* = 3.48e-02 and from 0.6 to 0.7 *p* = 4.02e-02; two-tailed cluster-based statistics). **b**. Time-resolved correlation between MI values and rhyming task accuracy. Notably, the correlation exhibits a temporal profile like that of neural encoding, though it peaks at earlier lags (dim 3: −0.2s lag; dim 7: 0s lag, when achieves statistical significance: *r* = 0.59, *p* = 3.51e-02, Bonferroni corrected). **c**. Normalized mean mutual information between CL latent and wav2vec 2.0 PCA subspaces for the model with 7 output dimensions. CL models encode PCA subspaces in a patter that is similar to what found in EEG encoding of the same subspaces. This pattern of bottom PCA subspaces encoding becomes more extreme as the CL models are forced to predict further in the future (from 20 to 200 ms ahead).

